# *Panax notoginseng saponins* protect the blood–brain barrier against oxidative stress by activating the Akap12–PI3K/AKT–AQP1 signaling axis

**DOI:** 10.1101/2025.11.14.688373

**Authors:** Lili Fan, Shukun Qu, Chenxue Miao, Dongjie Wu, Yuan Rao, Weisong Xu, Yanping He, Rui Ding, Guang Song

**Author notes:** Corresponding author: School of Life Sciences, Central China Normal University, NO.152 Luoyu Road, Wuhan 430079, China. E-mail addresses.

## Abstract

Oxidative stress is a key pathological factor that contributes to blood–brain barrier (BBB) disruption during ischemic injury. Panax notoginseng saponins (PNS), the primary bioactive components of Panax notoginseng, possess potent antioxidative and vasoprotective activities. This study aimed to elucidate the molecular mechanisms by which PNS regulate endothelial integrity and BBB permeability under oxidative stress conditions. Mouse brain microvascular endothelial cells (bEnd.3) were treated with hydrogen peroxide to induce oxidative damage, followed by intervention with PNS. Transcriptomic and proteomic analyses revealed that Akap12 (A-kinase anchoring protein 12) and Aquaporin 1 (AQP1) were key molecules involved in PNS-mediated endothelial protection. H₂O₂ exposure markedly suppressed Akap12 expression and inhibited the PI3K/AKT signaling pathway, leading to elevated AQP1 expression and increased BBB permeability. PNS treatment reversed these effects by restoring Akap12 expression, reactivating PI3K/AKT signaling, and reducing AQP1 levels. Functional validation further showed that AQP1 dysregulation altered the expression balance of tight junction proteins, including Claudin-5 and ZO-1, thereby impairing barrier function. Collectively, these findings demonstrate that PNS protect endothelial cells from oxidative injury by activating Akap12-dependent PI3K/AKT signaling and modulating AQP1-associated tight junction remodeling. This study provides new mechanistic insights into the molecular basis of PNS-mediated BBB protection and suggests potential therapeutic targets for oxidative or ischemic cerebrovascular diseases.

## 1. Introduction

The blood-brain barrier (BBB), recently redefined as the blood-brain boundary, serves a dynamic interface between the central nervous system (CNS) and the bloodstream. Disruption of the BBB exposes the CNS to peripheral pathogens and harmful substances, contributing to or exacerbating various CNS disorders [1]. Brain microvascular endothelial cells (BMECs), situated at the interface between brain tissue and blood, perform essential physiological functions, including barrier maintenance, nutrient transport, receptor-mediated signal transduction, and regulation of leukocyte infiltration[2]. As the primary barrier preventing blood components from entering the neural system, endothelial cells are the core components of the BBB, exhibiting unique structural and molecular characteristics. These include the expression of tight junction proteins that regulate the paracellular pathway and the expression of cell adhesion molecules[3]. During injury, endothelial cell activation triggers vascular remodeling, disrupts the tight junctions (TJs), and increases paracellular flux, thereby disrupting brain homeostasis and aggravating disease [4, 5]. Therefore, promoting vascular health and stability could be beneficial for slowing disease progression and supporting recovery. However, to date, our understanding of the spatiotemporal alterations of the BBB and the mechanistically regulated processes occurring during brain injury remains limited.

Hypoxia is implicated in various brain disorders[6]. At the cellular level, the adaptive response to hypoxia is mediated by signaling molecules such as reactive oxygen species (ROS)[7] and transcription factors, such as hypoxia-inducible factors (HIFs). Hypoxia can trigger or mitigate oxidative stress, mitochondrial dysfunction, and inflammation [8]. Modulating hypoxia offers a promising approach for treating CNS disorders. Hypoxia-induced disruption of BBB exacerbates cerebral stroke[9]. Additionally, HIFs have proved to be the regulatory roles in inflammatory bowel disease [10].

*Panax notoginseng* (Burkill) F.H. Chen, the dried root and rhizome of the plant, is a traditional Chinese medicine known as Sanqi or Tianqi in China, with a documented therapeutic history dating back to 1578 AD[11, 12] [13]. Advances in modern pharmacology have contributed to its widespread use in Chinese medicinal preparations, particularly due to its well-recognized efficacy in promoting blood circulation and resolving blood stasis[14]. The pharmacological activities of *Panax notoginseng* are primarily attributed to its bioactive components, collectively known as *Panax notoginseng* saponins (PNS). These include five major saponins: notoginsenoside R1 (NG-R1), ginsenoside Rg1 (G-Rg1), ginsenoside Re (G-Re), ginsenoside Rb1 (G-Rb1), and ginsenoside Rd (G-Rd), which are present in varying proportions [15]. Numerous studies have demonstrated that PNS exhibit antioxidant, anti-inflammatory, and other beneficial effects.

PNS have been shown to promote skin wound healing and inhibit scar formation in mice [16]. Furthermore, total saponins from *Panax notoginseng* exhibit therapeutic potential for diabetes-induced retinopathy [17]. PNS also protect against cerebral ischemia-reperfusion injury by reducing inflammation and modulating oxidative stress [1]. However, the molecular mechanisms by which PNS regulate hypoxia-induced alterations in BBB permeability remain poorly understood.

In this study, we employed integrated transcriptomic and proteomic analyses to investigate the molecular mechanisms through which PNS reduce hypoxia-induced alterations in BBB permeability. Our results demonstrate that PNS alleviates the hypoxia-induced increase in BBB permeability in vitro. Knockdown and overexpression experiments identified AQP1 as a critical factor influencing BBB permeability.

## 2. Materials and methods

### 2.1 Cell culture

bEnd.3 cells were purchased from Wuhan Pricella Biotechnology (CL-0598), bEnd.3 cells were cultured at 37℃ in a humidified atmosphere of 5% CO_2_ and 95% air in DMEM containing 10% heat-inactivated fetal bovine serum (BBI, Shanghai, China). bEnd.3 cells were treated with different concentrations of H_2_O_2_ (0, 200, 400, 600, 800 and 1000 µm) and PNS (0, 0.2, 0.4, 0.8, 1.2 and 1.6 mg/mL) for 24 h. Subsequently, the cells were incubated with 100 µL of DMEM and 10 µL of Cell Counting Kit-8 Reagent (Abbkine Scientific Co., Ltd, China) in the dark for 1h at 37°C. The optical density (OD) was measured at 450 nm using a Synergy H1 Reader (BioTek, Winooski, VT, USA).

### 2.2 Wound healing assay

For the horizontal migration assay, 3 × 10⁵ cells were seeded into six-well plates and cultured until reaching full confluence. A linear scratch was then created in each well using a sterile pipette tip, and the wells were washed twice with PBS to remove detached cells. Fresh DMEM containing 800 µM H₂O₂ was added, and the cells were incubated for 24 h. The wound closure was photographed at 0 and 24 h using an inverted microscope, and the migration rate was calculated using ImageJ software (version 1.41; NIH).

### 2.3 Transwell migration assay

For the vertical migration assay, 2 × 10⁴ cells were seeded into the upper Transwell chamber (8-µm pore size) containing serum-free DMEM, while the lower chamber was filled with DMEM supplemented with 20% FBS. After incubation for 24 h, non-migrated cells on the upper surface of the membrane were gently removed with a cotton swab. The migrated cells on the lower surface were fixed with 4% paraformaldehyde for 20 min and stained with 0.1% crystal violet for 15 min. Images were captured using the ImageView system at 10× magnification. The number of migrated cells was quantified using ImageJ and visualized with GraphPad Prism 9.0.

### 2.4 BBB permeability assay

The permeability of the BBB was evaluated using sodium fluorescein (NaFI) as a tracer. Briefly, 100 µL of NEST GelNest Matrix (LDEV-free) was added to the upper chamber of a Transwell insert and incubated at 37 °C with 5% CO₂ for 2 h. Subsequently, 2 × 10⁴ cells per well were seeded into the upper chamber, while the lower chamber was filled with DMEM containing 20% FBS. The 24-well plate was incubated at 37 °C with 5% CO₂ for 24 h to allow cell monolayer formation. After incubation, 100 µL of NaFI solution (100 µg/mL in PBS) was added to the upper chamber, and the medium in the lower chamber was replaced with PBS. Fluorescence intensity in the lower chamber was measured at 0, 15, 30, 60, and 90 min using a BioTek Synergy H1 microplate reader (excitation: 470 nm; emission: 520 nm).

A time-course plot of fluorescence was generated, and the slope of the linear regression was defined as Pₜ, representing total clearance. BBB permeability (Pₑ) was calculated according to the following formula: 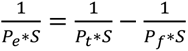, where *S* represents the surface area of the membrane, Pₜ is total permeability, and P_f_ is permeability without cells. The clearance volume (V_c_) was calculated as: *V*_e_ = (*V*_b_ ∗ *C*_b_)/*C*_t_. where *V_b_* is the volume of the lower chamber, *C_b_* is the concentration of NaFI in the lower chamber, and *C_t_* is the concentration in the upper chamber.

### 2.5 Measurement of reactive oxygen species (ROS)

Reactive oxygen species (ROS) were measured using 2’,7’-dichlorodihydrofluorescein diacetate (DCFH-DA, aladdin). bEnd.3 cells were washed with phosphate-buffered saline (PBS) and incubated in serum-free basal medium containing 10µM DCFH-DA for 20 min at 37°C. Then, the cells were washed twice with PBS. Fluorescence images were captured immediately using EVOS M7000 Imaging System (Thermo Fisher Scientific, CA, USA) with an excitation wavelength of 488 nm and an emission wavelength of 510 nm. The green fluorescence intensity of DCFH-DA in each cell was captured using the EVOS™ M7000 imaging system and quantified using ImageJ software (version 1.41; NIH).

### 2.6 L-LDH, SOD and CAT activity

The activities of the antioxidant enzymes catalase (CAT) and superoxide dismutase (SOD) were measured using the Catalase Assay Kit (Solarbio, Catalase(CAT)Activity Assay Kit, Cat:BC0205) and the Micro-method SOD Activity Detection Kit (Solarbio, Superoxide Dismutase(SOD)Activity Assay Kit(WST-1 Method), Cat: BC5165) respectively. Five million cells were mixed with 1 mL PBS or superoxide dismutase lysis buffer, lysed by ultrasonication and then centrifuged at 8000 g for 10 min at 4°C, and the supernatant was collected for analysis. The absorbance of CAT and SOD was measured at 560nm and 405 nm respectively, using Synergy H1 Reader (BioTek), and the result was expressed as Uꞏmg^−1^ protein. Lacate Dehydrogenase (LDH)Activity Assay Kit (Solarbio, Cat: BC0685) was used to measure the lactate dehydrogenase activity. cells were mixed with 1 mL LDH extract, sonicated (3 seconds on, 10 seconds off, repeated 30 times) and then centrifuged at 8,000 g at 4°C for 10 min. Then the supernatant was analyzed at 450nm using Synergy H1 Reader (BioTek), with results expressed as U/10^4^ cells.

### 2.7 Transcriptome sequencing and data analysis

Total RNA extraction, library construction, sequencing and were performed by Beijing Biomarker Biotechnology Co., Ltd., following their standard operating procedures. Briefly, total RNA was isolated using TRIzol reagent (Life Technologies, CA, USA) according to the manufacturer’s protocol. RNA concentration and purity were determined using a NanoDrop 2000 spectrophotometer (Thermo Fisher Scientific, DE, USA), and RNA integrity was evaluated with the RNA Nano 6000 Assay Kit on an Agilent 2100 Bioanalyzer (Agilent Technologies, CA, USA). Library quality was also assessed using the Agilent 2100 Bioanalyzer. The qualified libraries were then sequenced on an Illumina NovaSeq platform to generate 150 bp paired-end reads.

After removing the adapter sequences and low-quality reads were removed from the datasets. Raw sequences were transformed into clean reads after data processing. These clean reads were then mapped to the reference genome (Mus musculus, GCA_000001635.8, http://asia.ensembl.org/index.html). Only reads with a perfect match or one mismatch were further analyzed and annotated using the reference genome. TopHat2 [28] was used to map the reads to the reference genome. Cufflinks were then employed to assemble the mapped reads and compare them with the annotation of the reference genome to identify new transcripts. The resulting new unigenes were subjected to BLAST searches and annotated against the NR, Swiss-Prot, GO, COG, KOG, Pfam, and KEGG databases. Cuffdiff was then used to identify differentially expressed genes between two sample groups, with threshold criteria of FDR ≤ 0.01 and a log2 fold change (0 h/48 h samples) ≥ 2, categorizing transcripts as up-regulated and down-regulated. To elucidate the molecular mechanisms underlying oxidative stress and PNS-mediated repair, bioinformatic analyses were performed on the identified differentially expressed proteins (DEPs). The DEPs were annotated and classified into functional pathways using the Gene Ontology (GO, 2024 release) and Kyoto Encyclopedia of Genes and Genomes (KEGG, 2024 release) databases. Subsequently, Euclidean distance–based hierarchical clustering was applied to generate a heatmap illustrating the expression patterns of all significant DEPs. Finally, Gene Set Enrichment Analysis (GSEA) was conducted to identify significantly enriched biological processes and signaling pathways involved in oxidative stress responses and PNS-induced repair.

### 2.8 Data-independent acquisition (DIA) proteomics analysis

Cells were washed twice with PBS and lysed in RIPA lysis buffer supplemented with 1 mM phenylmethylsulfonyl fluoride (PMSF). The lysates were sonicated on ice (3 s on, 10 s off; 20% power) and clarified by centrifugation. Protein concentration was determined using the BCA Protein Assay Kit (catalog no. 23227; Thermo Fisher Scientific, Waltham, MA, USA).

For proteomic analysis, 100 µg of total protein from each group was reduced by adding dithiothreitol (DTT) to a final concentration of 10 mM, followed by incubation at 37°C with shaking at 180 rpm for 30 min. The reduced proteins were alkylated with iodoacetamide (IAM) at a final concentration of 15 mM and incubated at 37°C with shaking at 180 rpm in the dark for 30 min. The pH was adjusted to 7.5 prior to addition of trypsin at a protein-to-trypsin mass ratio of 50:1. Digestion was carried out at 37°C for 16 h and terminated by adding formic acid to a final concentration of 1%. Digests were centrifuged at 12,000 × g for 5 min, and the supernatant was collected. Peptides were desalted using a Strata X column and vacuum-dried using a CentriVap Refrigerated Concentrator (LABCONCO, Kansas City, MO, USA) prior to mass spectrometry analysis.

Mass spectrometry analysis was performed in data-independent acquisition (DIA) modes using nanoscale liquid chromatography–tandem mass spectrometry (nanoLC–MS/MS). All experiments were conducted on an UltiMate 3000 UHPLC system (Thermo Fisher Scientific, Waltham, MA, USA). The acquisition and analysis of DIA proteomics data are both performed by BGI Genomics (Shenzhen) Co., Ltd. same as previous published[18]. Differentially expressed proteins (DEPs) were set by a fold change >2 and *p* < 0.05. Bioinformatics analysis were performed as the previous analysis in the transcriptome data analysis.

### 2.9 Western blot analysis

Cells were lysed on ice with RIPA lysis buffer (Solarbio, Beijing, China). The lysates were centrifuged at 12,000 × g for 10 min at 4°C, and the resulting supernatant was collected as the total protein extract. Protein concentration was determined using the BCA Protein Assay Kit (Thermo Fisher Scientific, Waltham, MA, USA).

For Western blot analysis, 10 µg of total protein per lane was separated by SDS-PAGE (Bio-Rad, Hercules, CA, USA) and transferred to a PVDF membrane (Millipore, Billerica, MA, USA). Membranes were blocked with 5% non-fat dry milk in TBST and incubated overnight at 4°C with primary antibodies against AQP1, Claudin-5, ZO-1, AKT, GSK3β, and β-actin (all from Proteintech Group, Inc., Wuhan, China). After washing, membranes were incubated with horseradish peroxidase-conjugated secondary antibodies (Proteintech Group, Inc., Wuhan, China) for 1 h at room temperature. Protein bands were visualized using the ChemiDoc Imaging System (Bio-Rad Laboratories, Inc., Hercules, CA, USA) according to the manufacturer’s instructions. Band intensities were quantified using Image Lab software (Bio-Rad).

### 2.10 Cell Transfection

Cells were transfected with a negative control lentivirus (oe-NC) for 48 hours to serve as a baseline under hypoxic conditions. To overexpression of AQP1, cells were transfected with the plasmid pLV3-CMV-Aqp1 (mouse)-3×FLAG-CopGFP-Puro (Miaoling Biotech, Wuhan, China), in which the AQP1 coding sequence was designed based on the reference sequence from the NCBI database. While, for AQP1 knockdown, short hairpin RNA (shRNAs) targeting AQP1 were synthesized by Shengong Bio (Shanghai, China), and cloned into the pLV-EGFP(2A)-Puro. The sequences of the shRNA constructs are as follows:

shRNA-AQP1_1: Foreword oligo:5’-GATTCGCCACATTCTTCAGGTGCTTACTCGAGTAAGCACCTGAAGAATGTGGCTTTTTG-3’.

Reverse oligo: 5’-AATTCAAAAAGCCACATTCTTCAGGTGCTTACTCGAGTAAGCACCTGAAGAATGTGGC-3’

shRNA-AQP1_2: Foreword oligo:5’-GATTCGACAATTCACTTGGCCGCAATCTCGAGATTGCGGCCAAGTGAATTGTCTTTTTG-3’

Reverse oligo: 5’-AATTCAAAAA-GACAATTCACTTGGCCGCAAT-CTCGAG-ATTGCGGCCAAGTGAATTGTC -3’

### 2.11 Statistical analysis

The data are reported as mean ± standard deviation, derived from three independent studies. For the normally distributed data, one-way analysis of variance (ANOVA) followed by the Tukey test or two-way ANOVA followed by the Bonferroni multiple comparison test was conducted. for non-normally distributed data, the Kruskal-Wallis test was applied. Image J (version 1.41; NIH) and GraphPad Prism 9.0 (GraphPad Software Inc., San Diego, CA, USA) were used to perform the statistical analysis. Statistical significance was defined as *p* < 0.05.

## 3. Results

### 3.1 BBB Permeability increased under high oxidative stress

Previous studies have demonstrated that ischemic stroke disrupts the integrity and functionality of the BBB. To model the oxidative stress associated the post-ischemic injury in brain endothelial cells, we exposed bEnd.3 cells to ascalating concentrations of H₂O₂ (100, 200, 400, 600, and 800 μM) to determine the optimal dose for our experimental model. After 24 hours exposure to 800 μM H₂O₂, cell viability was significantly declined (**Fig. 1A and B**). Accordingly, 800 μM H₂O₂ was selected to establish an in vitro model of ischemic stroke.

**Fig. 1.**
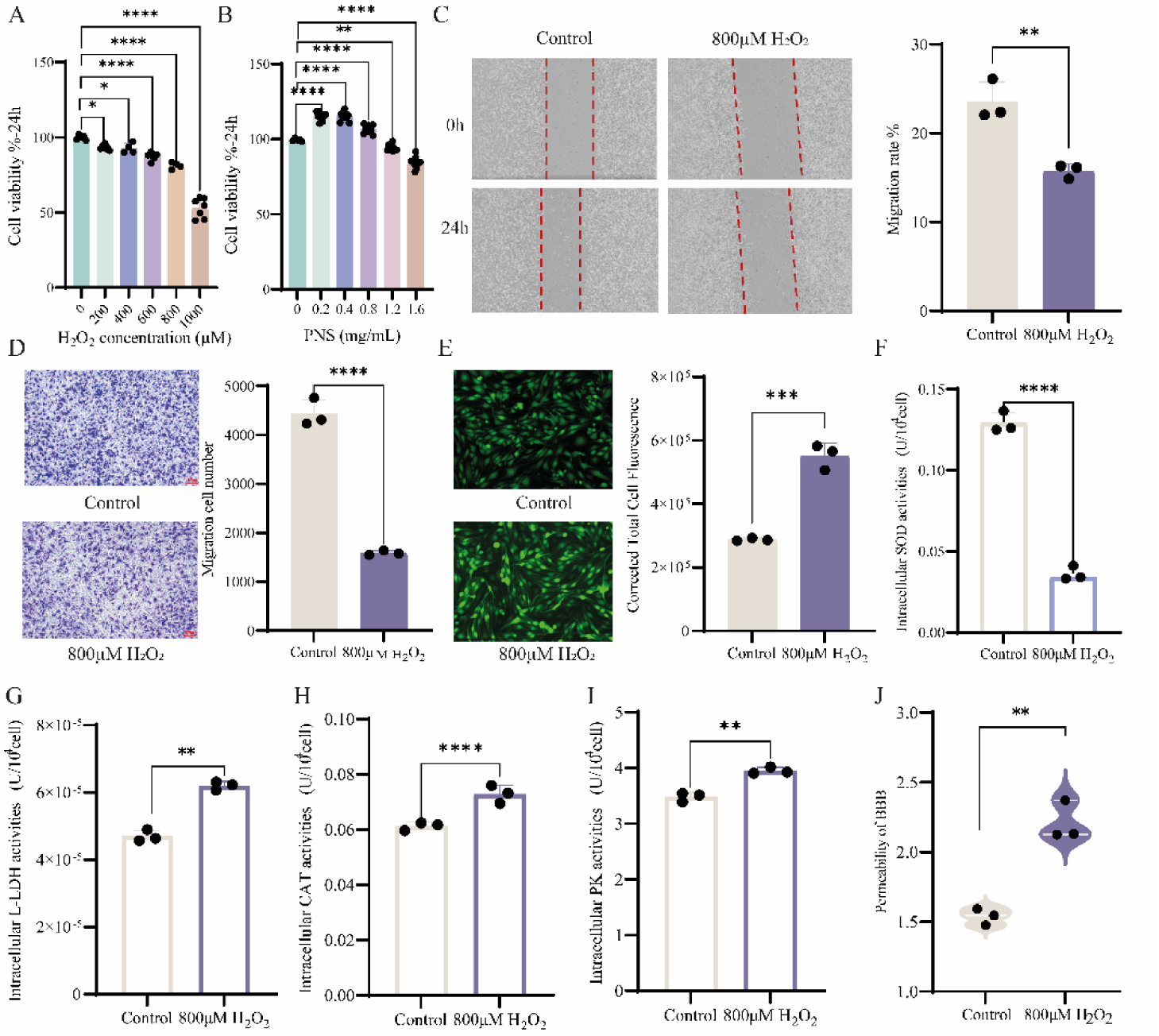
Effects of H₂O₂ on endothelial cell phenotype, enzyme activities, and BBB permeability. **(A)** Cell viability assessed by CCK-8 assay after treatment with different H₂O₂ concentrations; **(B)** Cell viability after H₂O₂ exposure followed by treatment with various concentrations of PNS; **(C)** Horizontal cell migration was evaluated using the wound-healing assay; **(D)** Vertical migration was examined by Transwell assay; **(E)** Intracellular ROS were visualized with 10 µM DCFH-DA using the EVOS™ M7000 system; **(F–I)** CAT, SOD, LDH, and PK activities were determined; **(J)** BBB permeability was measured using sodium fluorescein (NaFI). All data are shown as mean ± SD; \**p* < 0.05, \*\**p* < 0.01, \*\*\**p* < 0.001.

Scratch assays provide a robust method for assessing cellular processes underlying BBB formation and function. They offer insights into the migratory and proliferative capabilities of cells associated with BBB formation in the vicinity of a wound, thereby facilitating our understanding of the dynamic changes occurring during BBB development.

After treating cells with 800 μM H₂O₂ for 24 hours, we observed a significant reduction in migration rate compared to the controls (*p* < 0.01) **(Fig. 1C**). Similarly, H₂O₂ treatment markedly impaired transwell migration (*p* < 0.0001) (**Fig. 1D**).

DCFH-DA probe was employed to quantify the intracellular reactive oxygen species (ROS) levels post-H₂O₂ treatment, and the results indicated that 800 μM H₂O₂ treatment significantly elevated intracellular ROS levels (*p* < 0.001) (**Fig. 1E**). In light of this significant increase in ROS, we assessed the activities of enzymes linked with oxidative damage. As shown in Figure 1F-I, H₂O₂ treatment significantly decreased the activities of antioxidant enzymes, including catalase (CAT) and superoxide dismutase (SOD) (*p* < 0.001), as well as pyruvate kinase (PK) (*p* < 0.01) (**Fig. 1F-I**).

Post-stroke loss of endothelial cell integrity often leads to increased paracellular BBB permeability. To confirm the effect of H₂O₂ treatment on BBB permeability, we used sodium fluorescein dye as a tracer. Compared to the control groups, H₂O₂ treatment significantly enhanced BBB permeability (*p* < 0.01) (**Fig. 1J**). Together, these findings indicate that H₂O₂ treatment elevates BBB permeability and intracellular ROS levels, suggesting that intracellular oxidative stress modulates BBB functions.

### 3.2 PNS Attenuates H₂O₂-Induced Increase in BBB Permeability

PNS, the major active component of the traditional Chinese herb *Panax notoginseng*, has been reported to promote cell proliferation, reduce oxygen consumption, and enhance cellular tolerance to hypoxia. We treated cells with varying concentrations of PNS (0.2, 0.4, 0.8, 1.2, and 1.6 mg/mL) and observed that the proliferative effect of PNS increased with concentration up to an optimal level, beyond which higher concentrations exhibited cytotoxicity. The most pronounced promotion of cell proliferation occurred at a PNS concentration of 0.2 mg/mL (*p* < 0.05) (**Fig. 1A**).

Subsequently, we investigated whether PNS could mitigate H₂O₂-induced cell damage and alterations in BBB permeability. Scratch assays were performed to assess the impact of PNS on cell proliferation and migration following H₂O₂ treatment (**Fig. 2A**). The results demonstrated that 0.2 mg/mL PNS significantly enhanced the cell migration rate, which was otherwise reduced by H₂O₂ (*p* < 0.0001). Additionally, Transwell assays indicated that PNS significantly promoted cell migration (*p* < 0.0001) (**Fig. 2B**).

**Fig. 2.**
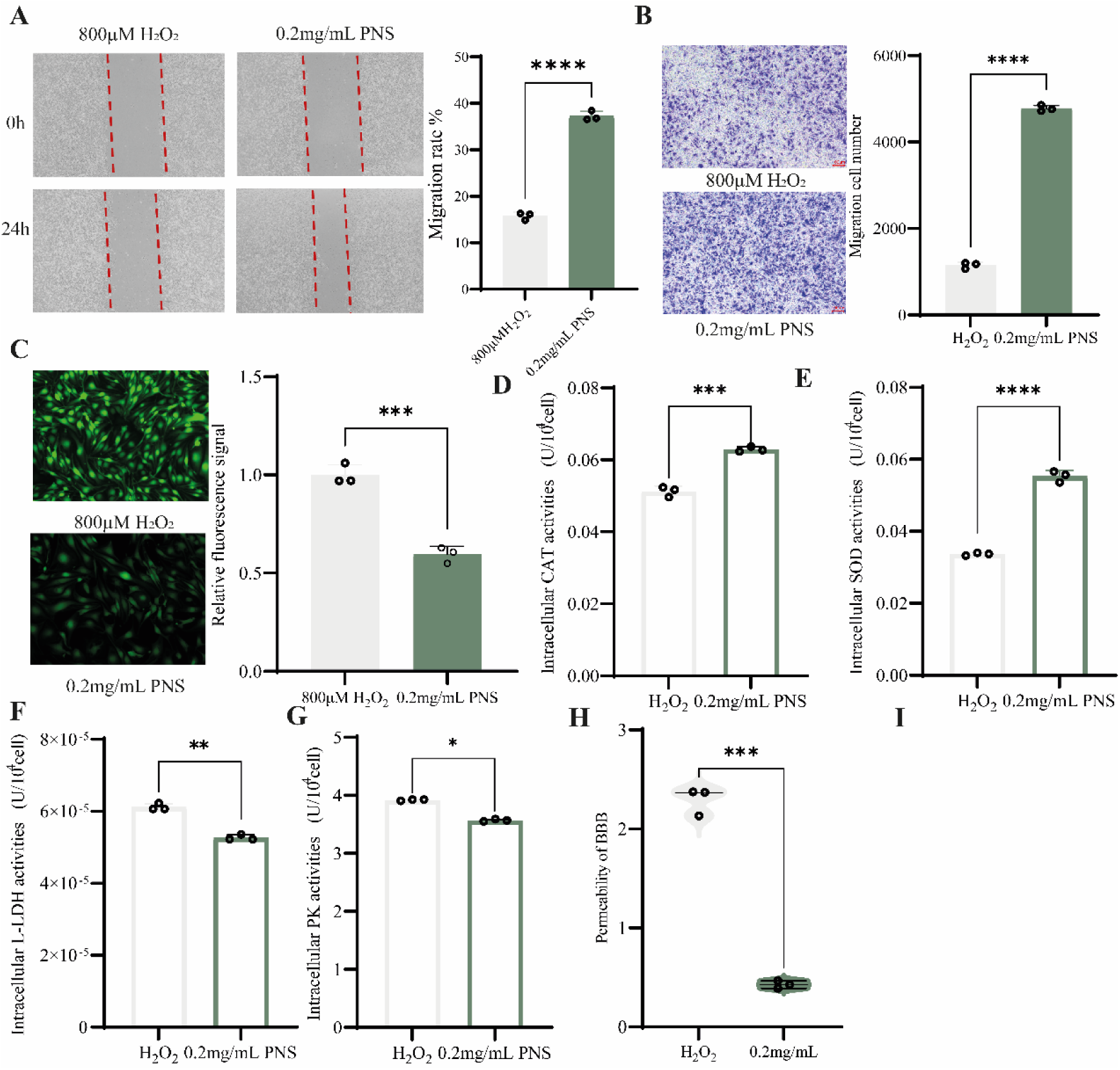
Effects of PNS on endothelial repair and BBB permeability after oxidative injury. **(A)** Horizontal migration was analyzed by wound-healing assay following PNS treatment after H₂O₂ exposure; **(B)** Vertical migration was evaluated by Transwell assay; **(C)** Intracellular ROS levels were measured; **(D–G)** Activities of CAT, SOD, LDH, and PK were determined using standard biochemical kits; **(H)** BBB permeability was determined using the NaFI assay; All data are expressed as mean ± SD; \**p* < 0.05, \*\**p* < 0.01, \*\*\**p* < 0.001.

To assess whether PNS could alleviate the H₂O₂-induced elevation in ROS levels, we utilized the DCFH-DA probe to quantify the intracellular ROS. The results indicated that, compared to H₂O₂-treated group, PNS significantly reduced the ROS induced by oxidative damage, as indicated by a notable reduction in fluorescence intensity (*p* < 0.001) (**Fig. 2C**).

Furthermore, to examine the effect of PNS on the activities of intracellular enzyme associated with oxidative stress following H₂O₂ treatment, we measured enzyme activities as a marker of intracellular oxidative damage. Consistent with our hypothesis, PNS treatment significantly elevated the activities of CAT, SOD, and PK enzymes compared to the H₂O₂-treated group. Moreover, PNS reversed the H₂O₂-induced increase in lactate dehydrogenase (LDH) activity (*p* < 0.05) (**Fig. 2D-G**).

Finally, we assess BBB permeability using sodium fluorescein tracing (**Fig. 2H**). Consistent with the previous findings, PNS treatment significantly attenuates the increases in BBB permeability induced by H₂O₂ treatment (*p* < 0.001). Collectively, these results suggest that PNS protects against H₂O₂-induced increases in BBB permeability by mitigating intracellular ROS levels.

### 3.3 Akap12-Dependent PI3K/AKT Activation in PNS-Mediated BBB Protection

Ischemia is well known to disrupt the BBB integrity [19]. There are two major cellular mechanisms underlie BBB dysfunction following ischemic stroke: Enhanced paracellular permeability due to disruption of endothelial cell tight junctions and increased transcellular permeability resulting from caveolae upregulation [3].To elucidate the molecular mechanisms regulating BBB permeability under oxidative stress in vitro, we performed transcriptomic profiling of mouse brain microvascular endothelial cells (bEnd.3) under three conditions: control, H₂O₂-treated, and PNS-treated at 24 h post H₂O₂ exposure and PNS intervention.

Compared with the control group, H₂O₂ treatment resulted in 2,411 downregulated and 1,933 upregulated genes (**sFig. 1 A and B**). Among the most significantly downregulated transcripts were Grhl2, Npnt, Mpp3, Nwd1, Akap12, Egr2, Egr1, and Adamts5. The functional enrichment analysis of these differentially expressed genes (DEGs) revealed significant enrichment of oxidative stress–related pathways (**sFig. 1C and D**), while Gene Set Enrichment Analysis (GSEA) indicated potential involvement in angiogenesis (**sFig. 1E**).

Upon PNS treatment, compared to the H_2_O_2_ - treated group, 208 genes were downregulated and 142 genes were upregulated (**sFig. 2A, and B**). The enrichment analysis of these DEGs again highlighted oxidative stress–associated pathways (**sFig. 2C and D**). whereas GSEA suggested a strong association with vascular morphogenesis (**sFig. 2E**).

To identify genes potentially mediating the reparative effects of PNS, we intersected the genes downregulated by H₂O₂ and upregulated by PNS, obtaining 226 overlapping candidates. Classification of these intersection genes revealed Akap12 (A-kinase anchoring protein 12) as a key molecule involved in signal localization, scaffolding, and the integration of PKA/Ca²⁺ signaling (**Fig. 3A**). KEGG and Gene Ontology (GO) analyses of the intersecting gene set showed significant enrichment in oxidative stress–related pathways, notably including the PI3K/AKT signaling pathway (*p* < 0.05), as well as the MAPK and calcium ion signaling pathways (**Fig. 3B and C**).

**Fig. 3.**
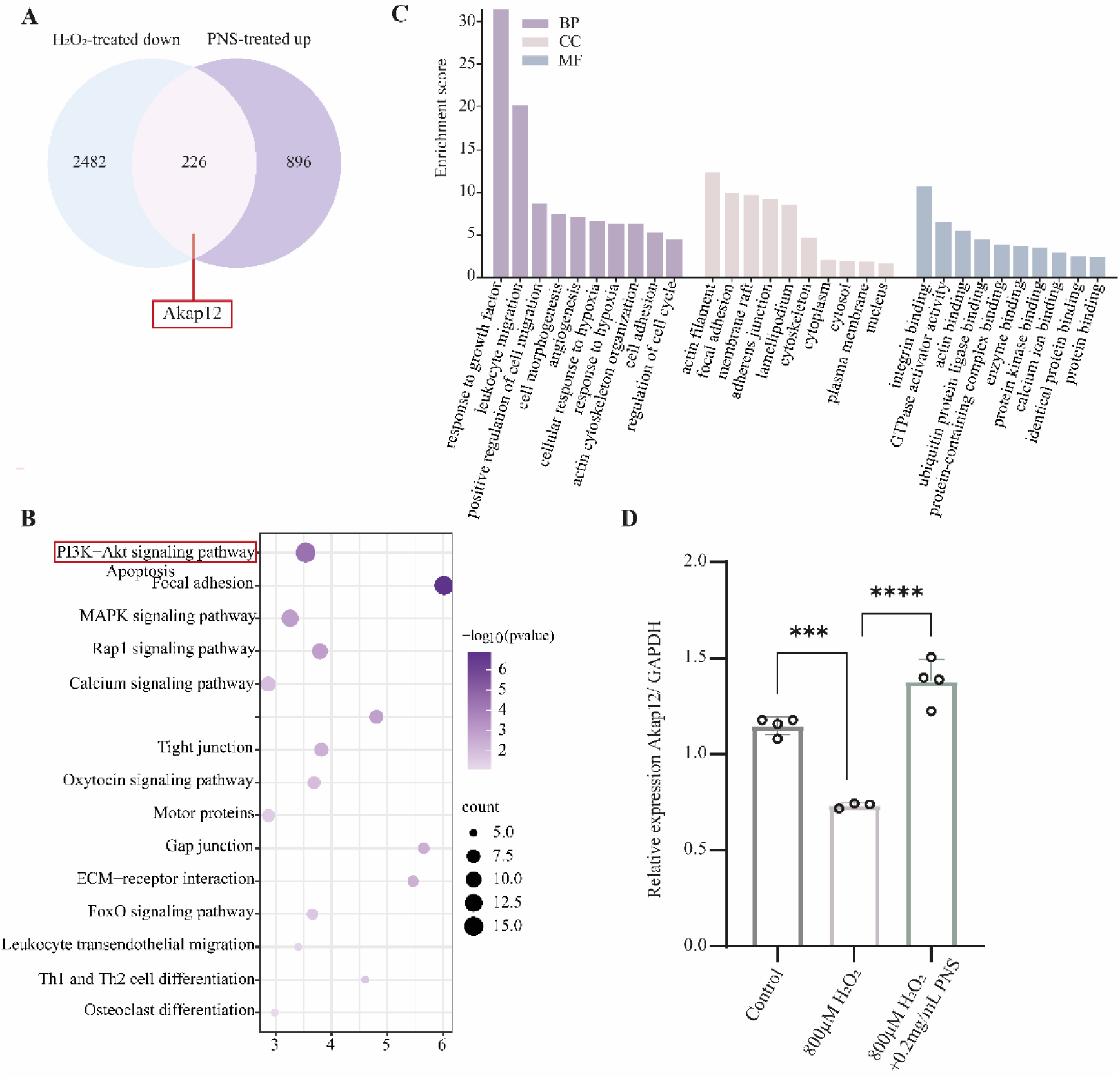
Transcriptomic profiling of H₂O₂-treated and PNS-treated endothelial cells. **(A)** Venn diagram showing counter-regulated genes which were downregulated by H₂O₂ and upregulated by PNS treatment. **(B)** Top 15 significantly KEGG pathways enriched in those counter-regulated genes; **(C)** Top 10 Gene Ontology (GO) terms for biological process (BP), cellular component (CC), and molecular function (MF) categories among the counter-regulated genes. **(D)** Quantitative PCR validation of Akap12 mRNA expression in control, H₂O₂-treated and PNS-treated cells. Data are shown as mean ± SD; \**p* < 0.05, \*\**p* < 0.01, \*\*\**p* < 0.001.

Quantitative real-time PCR (qPCR) analysis confirmed that Akap12 expression exhibited the same trend as observed in the transcriptomic data (**sFig. 1F and 2F**). Collectively, these findings suggest that Akap12 may serve as a signaling scaffold that bridges oxidative stress responses and PI3K/AKT pathway activation, mediating the protective effects of PNS against H₂O₂-induced endothelial injury.

### 3.4 PNS Downregulates AQP1 Expression to Ameliorate Endothelial Injury

To identify key proteins that play pivotal roles in endothelial injury and repair, we performed a comprehensive proteomic analysis to examine changes induced by H₂O₂ exposure and subsequent PNS treatment. After H₂O₂ stimulation, 1,263 proteins were significantly upregulated and 2,475 were downregulated (**sFig. 3A and B**). Functional enrichment analysis revealed that the upregulated proteins were predominantly involved in oxidative phosphorylation and phosphatidylinositol signaling pathways (**sFig. 3C and D**), suggesting enhanced oxidative metabolism and stress signaling.

Following PNS intervention, 1,274 proteins were upregulated and 2,464 were downregulated relative to the H₂O₂-treated group (**sFig. 4A and B**). These downregulated proteins were mainly enriched in the HIF1α and PI3K/AKT signaling pathways (**sFig. 4C and D**), indicating that PNS may suppress hypoxia- and stress-related signaling.

To further identify key mediators of PNS repair, we intersected the proteins upregulated by H₂O₂ with those downregulated by PNS treatment, yielding 202 overlapping candidates (**Fig. 4A**). Enrichment analysis of these overlapping proteins showed significant involvement in the PI3K/AKT, MAPK, and mTOR signaling pathways (**Fig. 4B-C**), all of which are crucial for endothelial survival and stress adaptation.

**Fig. 4.**
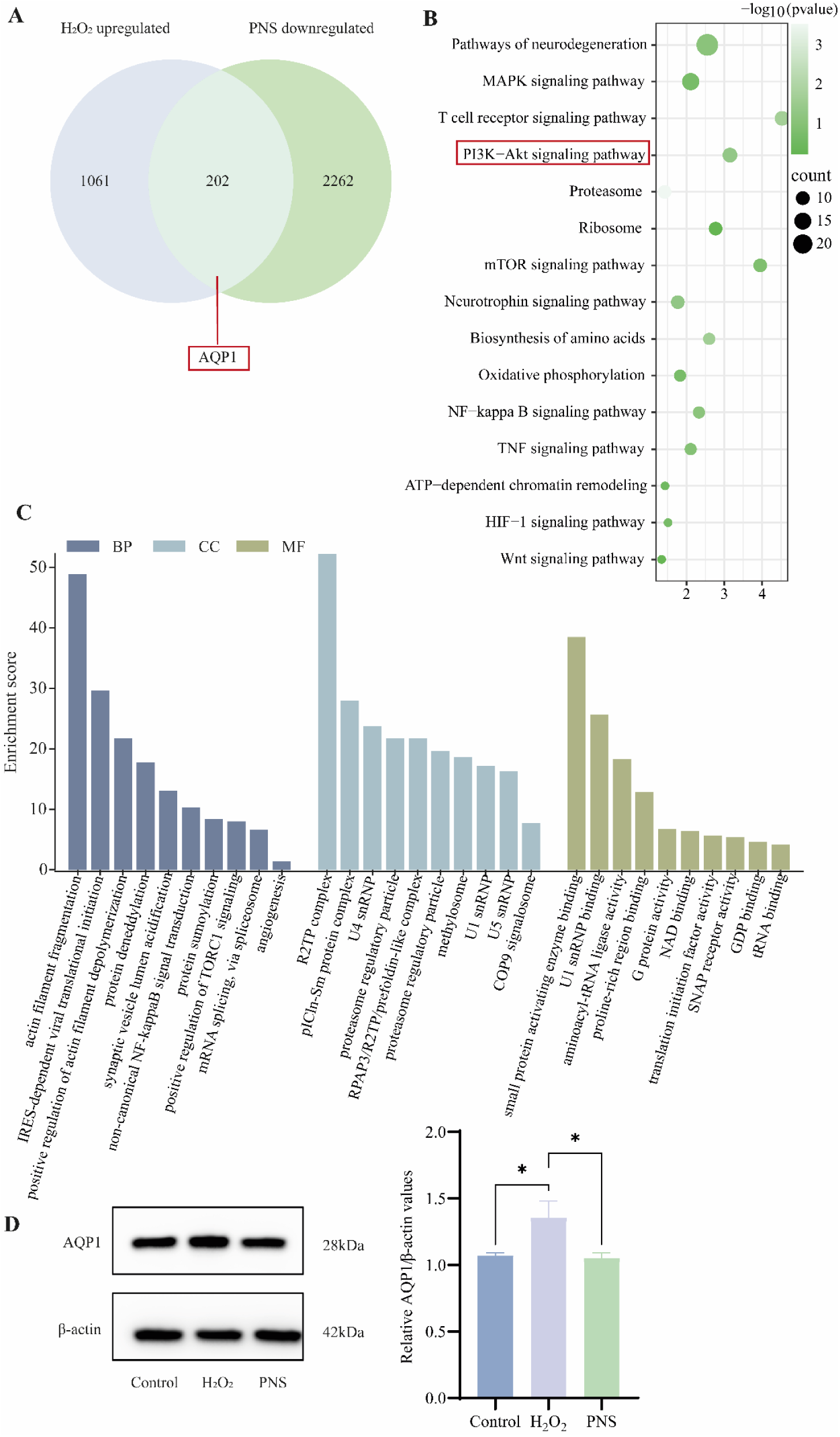
Proteomic analysis of control, H₂O₂-treated, and PNS-treated endothelial cells. **(A)** Volcano plot showing rescued proteins downregulated by H₂O₂ treatment and restored by PNS; **(B)** Top 15 significantly enriched KEGG pathways identified among those restored proteins; **(C)** GO functional classification showing the top 10 terms for BP, CC, and MF categories. **(D)** Relative protein expression of AQP1 normalized to β-actin was determined by western blot analysis. \**p* < 0.05.

Among these, Aquaporin 1 (AQP1) emerged as a notable candidate, exhibiting a marked increase in expression following H₂O₂ treatment and a significant decrease after PNS intervention. Western blot analysis confirmed these proteomic findings (**Fig. 4D**). Based on these results, we propose that PNS may attenuate endothelial injury by modulating AQP1 expression through Akap12-mediated PI3K/AKT signaling, thereby contributing to the restoration of endothelial function under oxidative stress.

### 3.5 PNS Reverses H₂O₂-Induced Suppression of the PI3K/AKT Pathway and Modulates AQP1 Expression

To further elucidate the involvement of the PI3K/AKT signaling pathway in oxidative damage and PNS-mediated repair, we employed the PI3K/AKT inhibitor LY294002 to assess how H₂O₂ and PNS influence the activation state of this pathway.

The wound-healing assay revealed that both H₂O₂ treatment and LY294002 treatment markedly suppressed endothelial cell migration compared with the control group (*p*<0.0001) (**Fig. 5A, B and C**). Consistently, reactive oxygen species (ROS) detection showed that both treatments led to a significant elevation in intracellular ROS levels (*p*<0.05) (**Fig. 5D–E**), suggesting comparable oxidative stress induction.

**Fig. 5.**
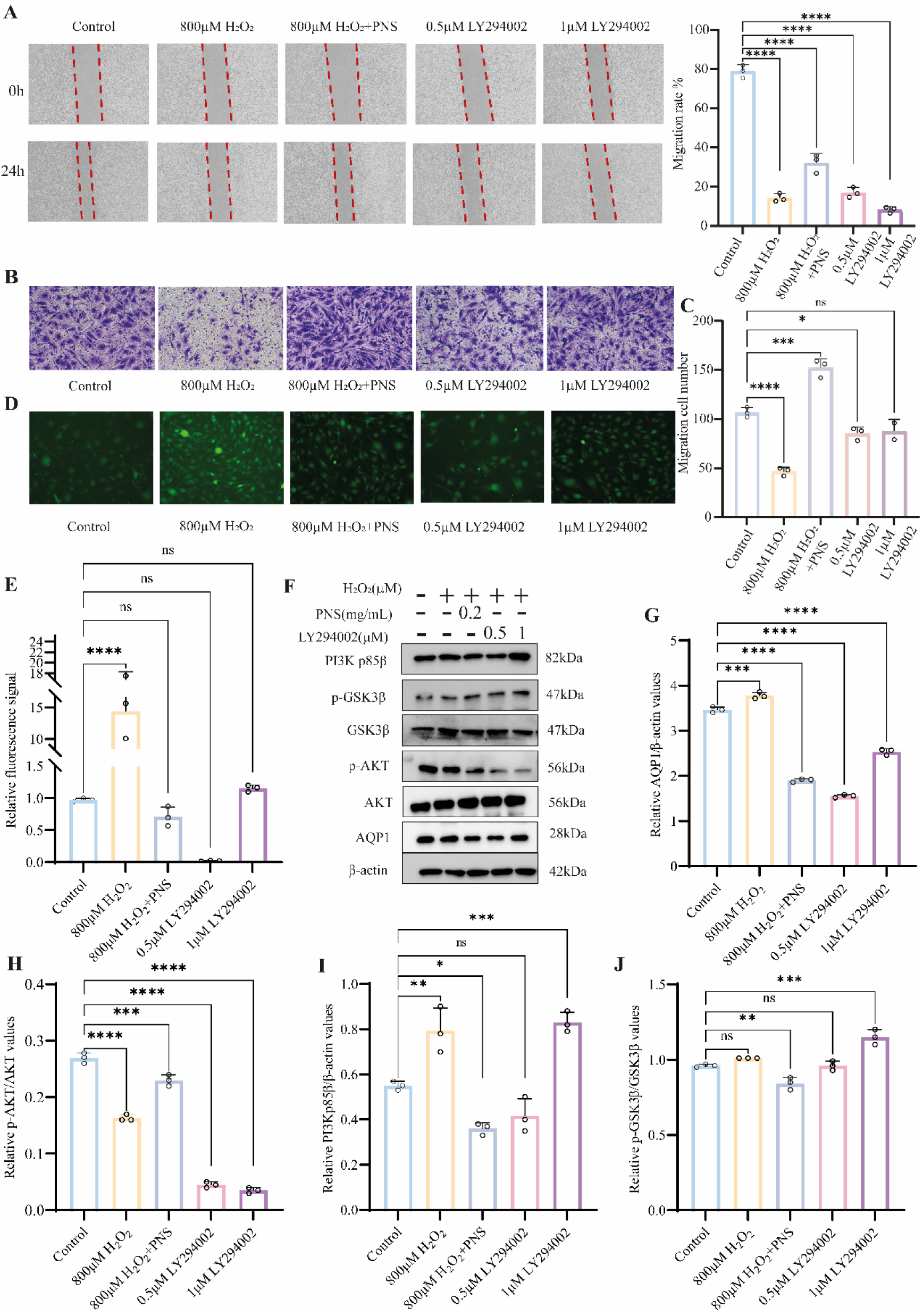
Inhibition of PI3K/AKT signaling by LY294002 and its effects on oxidative stress and AQP1 expression. **(A)** Horizontal cell migration analyzed by wound-healing assay; **(B–C)** Vertical migration assessed by Transwell assay; **(D–E)** ROS fluorescence measured by DCFH-DA probe and analyzed quantitatively; **(F)** Western blot analysis of β-actin, AQP1, AKT, p-AKT, GSK3β, p-GSK3β, and PI3K p85β; **(G–J)** Quantitative analysis of AQP1/β-actin, p-AKT/AKT, PI3K p85β/β-actin, and p-GSK3β/GSK3β ratios. Statistical significance was determined by one-way ANOVA. \**p* < 0.05, \*\**p* < 0.01, \*\*\**p* < 0.001.

Additionally, western blot analysis was then performed to examine the expression of key components of the PI3K/AKT pathway (**Fig. 5F–J**). Both H₂O₂ and LY294002 significantly reduced the phosphorylation level of AKT (p-AKT), indicating that oxidative stress inhibits the activation of the PI3K/AKT signaling pathway. Notably, inhibition of the PI3K/AKT pathway by LY294002 resulted in a significant reduction of AQP1 expression (*p* < 0.001), similar with the results with PNS treatment, implying that AQP1 is downstream of this pathway. These results suggest that PNS may affect AQP1 expression and thereby influence BBB permeability by reversing the inhibition of the PI3K/AKT pathway.

### 3.6 AQP1 Regulates BBB Permeability by Modulating Claudin-5 Expression

Transcriptomic and proteomic analyses suggested that oxidative stress induced by H₂O₂ disrupted angiogenic and vascular repair processes involving endothelial cells, with Aquaporin 1 (AQP1) emerging as a potential key regulator. To further investigate how AQP1 influences BBB permeability, we manipulated its expression in bEnd.3 cells using a lentiviral packaging system to achieve either AQP1 knockout or overexpression.

The wound healing assay revealed that AQP1 knockout significantly reduced the horizontal migration rate of endothelial cells (*p* < 0.05), whereas AQP1 overexpression markedly enhanced cell migration (*p* < 0.01) (**Fig. 6A**). A similar trend was observed in vertical migration assays (**Fig. 6B and C**), indicating that AQP1 facilitates endothelial motility.

**Fig. 6.**
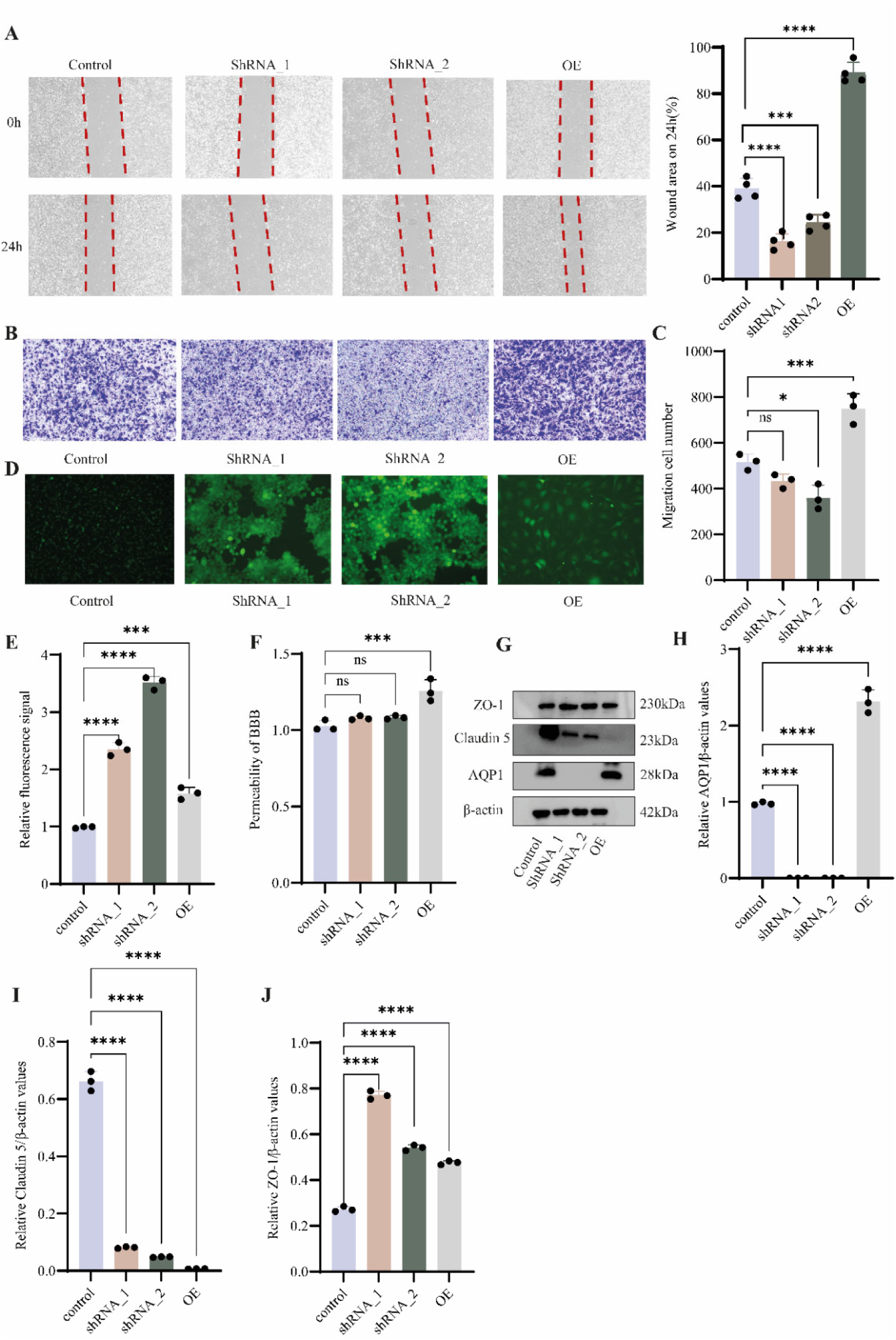
Knockout or overexpression of AQP1 alters endothelial migration and BBB permeability. **(A)** Horizontal migration rate determined by wound-healing assay after AQP1 knockout or overexpression. **(B–C)** Vertical migration evaluated by Transwell assay; migrated cells were visualized at 10× magnification and quantified. **(D–E)** Intracellular ROS levels detected with DCFH-DA; **(F)** BBB permeability determined using NaFI assay; **(G)** Western blot analysis of β-actin, AQP1, Claudin-5, and ZO-1; **(H–J)** Relative protein expression of AQP1/β-actin, ZO-1/β-actin, and Claudin-5/β-actin. Data analyzed by one-way ANOVA; \**p* < 0.05, \*\**p* < 0.01, \*\*\**p* < 0.001.

Next, intracellular reactive oxygen species (ROS) levels were measured using a DCFH-DA probe. Both AQP1 knockout and overexpression significantly increased ROS accumulation (*p* < 0.01) (**Fig. 6D and E**), suggesting that AQP1 dysregulation perturbs cellular redox balance. To assess the impact of AQP1 on BBB integrity, sodium fluorescein permeability assays were performed. The results showed that AQP1 overexpression led to increased BBB permeability (**Fig. 6F**).

Western blot analysis further demonstrated that both AQP1 knockout and overexpression significantly reduced the expression of the tight junction protein claudin-5, while concomitantly increasing the expression of another tight junction protein, ZO-1(*p*<0.001) (**Fig. 6G–J**). These results suggest that aberrant AQP1 expression disrupts the homeostatic balance between tight junction components, leading to altered junctional organization and increased BBB permeability.

Collectively, these findings indicate that AQP1 modulates BBB permeability by differentially regulating the expression of claudin-5 and ZO-1, thereby influencing tight junction remodeling and endothelial barrier integrity under oxidative stress conditions.

## 4. Discussion

Hypoxia is a persistent condition during both the initial and reparative phases of cardiovascular and cerebrovascular injury. Sustained hypoxia disrupts tissue homeostasis and repair, leading to metabolic stress, the accumulation of reactive oxygen species (ROS), and activation of autophagy [20]. Elevated ROS levels under hypoxic conditions contribute to oxidative stress, resulting in the oxidation of DNA, lipids, and proteins. Oxidative DNA damage can generate mutagenic adducts such as 8-oxo-7,8-dihydro-2′-deoxyguanosine (8-oxo-dG), a hallmark of hydroxyl radical attack on guanine residues [21].

Hypoxia exerts profound regulatory effects on protein expression involved in energy metabolism, angiogenesis, erythropoiesis, and autophagy[22] [6, 21]. Prolonged exposure to hypoxia impairs vascular development, alters blood circulation, and compromises the selective permeability of the blood–brain barrier (BBB), thereby promoting neuroinflammation and tissue edema [23, 24].

Previous studies have demonstrated that Aquaporin-1 (AQP1) expression is upregulated under hypoxic conditions through hypoxia-inducible factor-1α (HIF-1α)-dependent transcriptional activation[25]. The HIF-1α–AQP1 axis has been implicated in edema formation following nerve injury and myocardial infarction [26, 27]. BBB disruption and subsequent edema formation are well-recognized hallmarks of ischemic stroke [2, 24, 28, 29]. Interestingly, AKAP12 expression has been reported to inversely correlate with post-stroke edema severity, suggesting its protective role in maintaining BBB integrity[28, 30–32].

AKAP12 acts as a scaffold protein and negative mitogenic regulator, orchestrating cytoskeletal dynamics, endothelial stability, and barrier function in both the brain and retina. Through its scaffolding domains, AKAP12 coordinates PKA/Ca²⁺-dependent signaling and modulates downstream effectors, including the PI3K/AKT pathway [31, 33, 34]. The PI3K/AKT signaling cascade is a central survival pathway that mitigates oxidative damage, regulates angiogenesis, and preserves BBB integrity under ischemic conditions [19, 35, 36]. Pharmacological activation of this pathway has been shown to attenuate ischemia-induced BBB leakage and neuronal apoptosis [20, 26, 29, 37–41].

In the present study, we observed that H₂O₂-induced oxidative stress significantly downregulated Akap12 expression in brain microvascular endothelial cells (bEnd.3), coinciding with inhibition of PI3K/AKT signaling. Akap12 functions as a signaling platform that recruits and activates PI3K, thereby facilitating AKT membrane localization and phosphorylation. This activation triggers a cascade of downstream anti-apoptotic and metabolic effects that enhance cell survival under oxidative stress. Reduction of Akap12 expression disrupts this protective signaling, potentially compromising endothelial integrity and increasing BBB permeability.

Our data further demonstrate that treatment with Panax notoginseng saponins (PNS) reverses H₂O₂-induced suppression of Akap12 and restores PI3K/AKT pathway activation. This signaling restoration was accompanied by reduced AQP1 expression and improved BBB stability. Mechanistically, AQP1 dysregulation alters the expression balance of tight junction proteins—decreasing claudin-5 while increasing ZO-1—which collectively disrupts tight junction homeostasis and elevates BBB permeability. PNS intervention normalized these changes, suggesting that PNS exerts its protective effects by activating the Akap12–PI3K/AKT–AQP1 axis, thereby maintaining tight junction integrity and BBB function under oxidative stress conditions.

Taken together, our findings propose a mechanistic model in which PNS promotes endothelial repair and barrier protection through Akap12-mediated activation of PI3K/AKT signaling. This activation suppresses aberrant AQP1 expression, stabilizes tight junction protein composition, and mitigates oxidative stress–induced BBB dysfunction. These results provide new insights into the molecular basis of PNS-mediated cerebrovascular protection and highlight Akap12 and AQP1 as potential therapeutic targets for ischemic or oxidative vascular injury.

## CRediT authorship contribution statement

Lili Fan: Writing –original draft, Data curation, Visualization, Validation, Software, Methodology, Investigation, Formal analysis, Conceptualization; Shukun Qu: Software, Visualization. Chenxue Miao: Investigation, Visualization. Dongjie Wu: Methodology, Data curation; Yuan Rao: Methodology, Data curation; Weisong Xu: Methodology, Data curation; Yanping He: Resources; Rui Ding: Resources; Guang Song: Conceptualization, Supervision, Original draft, Writing-review & editing, Funding acquisition.

## Acknowledgments

This work was supported by the Fundamental Research Funds for the Central Universities (Grant No.CCNU25JC035) and Hubei Key Laboratory of Genetic Regulation and Integrative Biology (GRIB202507).

## Data availability

Any data can be sought from the corresponding author.

